# Reversible expression of a cartilage ‘master regulator’ during axolotl limb development and regeneration

**DOI:** 10.64898/2026.11.10.765670

**Authors:** Miray N. Bozkurt, Leo Otsuki, Elly M. Tanaka

**Author notes:** Corresponding Authors: LO, EMT.

## Abstract

How the expression dynamics of cell fate transcription factors relate to emerging cell identities is not well understood. The axolotl limb is a powerful model for investigating the acquisition of cell fate: during development, the limb mesenchyme differentiates into multiple cell types; during regeneration, missing cell types are reconstituted. Here, we investigate the expression of SOX9, a transcription factor widely regarded as a ‘master regulator’ of cartilage cell fate. By genetically fate mapping *Sox9*-expressing cells during development, we found that they give rise to two populations: cells that retain *Sox9* expression and become cartilage or perichondrial cells, and cells that extinguish *Sox9* expression and acquire alternative identities, such as dermal or interstitial fibroblasts. These populations differ in their contributions to regeneration: cells that retained *Sox9* expression contribute little to skeletal regeneration, while cells that downregulated *Sox9* contribute extensively, can reinitiate *Sox9* expression and form new cartilage. We found that transiently *Sox9*-expressing cells are also present during early limb regeneration. Together, our findings link sustained or transient expression of a cartilage ‘master regulator’ to distinct cell fates and subsequent contributions to skeletal regeneration.

## Introduction

Understanding how diverse cells contribute towards tissue formation is a fundamental question in developmental and regenerative biology. A foundational model to study the acquisition of cell fate has been the developing vertebrate limb. The early limb bud mesenchyme contains multipotent progenitors that have the potential to generate lateral plate mesoderm-derived cell types and tissues including cartilage, tendons, perichondrium, and fibroblasts (Arques et al., 2007; Pearse et al., 2007). As limb development proceeds, a subset of these progenitors initiates expression of the cartilage ‘master regulator’ SOX9, a transcription factor that promotes differentiation into a cartilage identity and whose expression is associated with cartilage tissue. In humans, heterozygous loss-of-function mutations in *SOX9* can lead to campomelic dysplasia, a severe skeleton malformation syndrome (Foster et al., 1994; Wagner et al., 1994). Chimera experiments in mice showed that *Sox9*−/− cells do not contribute to cartilage formation and lack expression of chondrocyte-specific markers (Bi et al., 1999). Conditional *Sox9* knockout resulted in lack of mesenchymal condensation or generalized chondrodysplasia with shorter skeletal structures and defective differentiation, depending on whether deletion was performed before or after the onset of chondrogenic condensation (Akiyama et al., 2002). Conversely, *Sox9* misexpression induced polydactyly characterized by ectopic *Sox5*- and *Sox6*-expressing condensations (Akiyama et al., 2007). Thus, *Sox9* is strongly linked to cartilage formation in the limb.

Given its functional importance, studying the expression of *Sox9* in relation to nascent cell identity and cartilage formation may contribute to an understanding of how cell identity factors control cell fate. Akiyama and colleagues previously performed genetic fate mapping of *Sox9*- expressing cells in the mouse limb bud (Akiyama et al., 2005). At E13.5, *Sox9*-expressing cells were present in the cartilage – as expected from *Sox9*’s role as a chondrogenic factor. By contrast, when *Sox9*-expressing cells were traced throughout limb development, these cells were found not only in cartilage but also in ligaments, tendons and synovia. A likely interpretation is that *Sox9* is initially expressed in a wider cell population than cells committed to cartilage differentiation, suggesting that *Sox9* alone does not irreversibly commit cells to cartilage identity. Yet the broader conservation of this expression pattern in other species, and its potential implications for cell function, have not been investigated.

In this study, we determined the dynamic features of *Sox9* expression in relation to emerging limb progenitor identities in the axolotl (*Ambystoma mexicanum*). Axolotls are four-legged vertebrates that exhibit overall conserved limb structure and genetic regulation compared with mice, humans and other vertebrates, with some regulatory differences (Christensen et al., 2002; Nowoshilow et al., 2018; Schloissnig et al., 2021). Axolotls can fully regenerate their limbs after injury, providing a unique opportunity to study cell fate decisions: first during limb development, then during limb regeneration. Amputation of an axolotl limb induces a proliferative mass of cells, called a blastema, to accumulate at the injury plane. Cells in the blastema differentiate to restore the missing cell types. Interestingly, skeleton is not regenerated by pre-existing skeletal cells in this injury paradigm (Currie et al., 2016; Gerber et al., 2018; McCusker et al., 2016).

Instead, the blastema contains multipotent mesenchymal cells that originate from loose fibroblastic cells in the limb: these express the transcription factor *Prrx1* and regenerate cartilage, tendons, ligaments, and fibroblasts, similar to mesenchymal cells in mouse or chick limb buds (Currie et al., 2016; Dunis and Namenwirth, 1977; Gerber et al., 2018; Hirata et al., 2010; Kragl et al., 2009; McCusker et al., 2016; Muneoka et al., 1986; Pearse et al., 2007). As in other vertebrates, both nascent and differentiated cartilage express *Sox9*. However, the dynamics of *Sox9* expression are not well understood, and genetic fate mapping has not been performed.

By carrying out time-resolved genetic fate mapping in axolotls, we identified *Sox9*-expressing cells during early limb development or regeneration that later lose *Sox9* expression and become non-cartilage cell types such as dermal or interstitial fibroblasts. Unlike cartilage, these cells can contribute extensively to limb regeneration, re-initiate *Sox9* expression and produce new cartilage. Thus, on-to-off *Sox9* expression is a conserved feature of mouse and axolotl limb development and also occurs during axolotl limb regeneration. Cells that switch off *Sox9* during development display greater contributions to limb regeneration than cells that retain *Sox9* expression.

## Results

### *Sox9* knock-in transgenic axolotls faithfully report *Sox9* expression

To study the spatiotemporal features of *Sox9* expression and relate these to cell identity in the limb bud, we made use of our recently reported *Sox9* transgenic axolotl line that co-expresses nuclear-localized EGFP and tamoxifen-inducible Cre recombinase from the endogenous *Sox9* gene locus (Tardivo and Olmeda et al., 2026) (Figure 1A). We expect this reporter to convert cells expressing SOX9 protein, as EGFP fluorescence requires translation of upstream *Sox9* and ribosome skipping at a T2A sequence (De Felipe et al., 2003; Donnelly et al., 2001).

**Figure 1.**
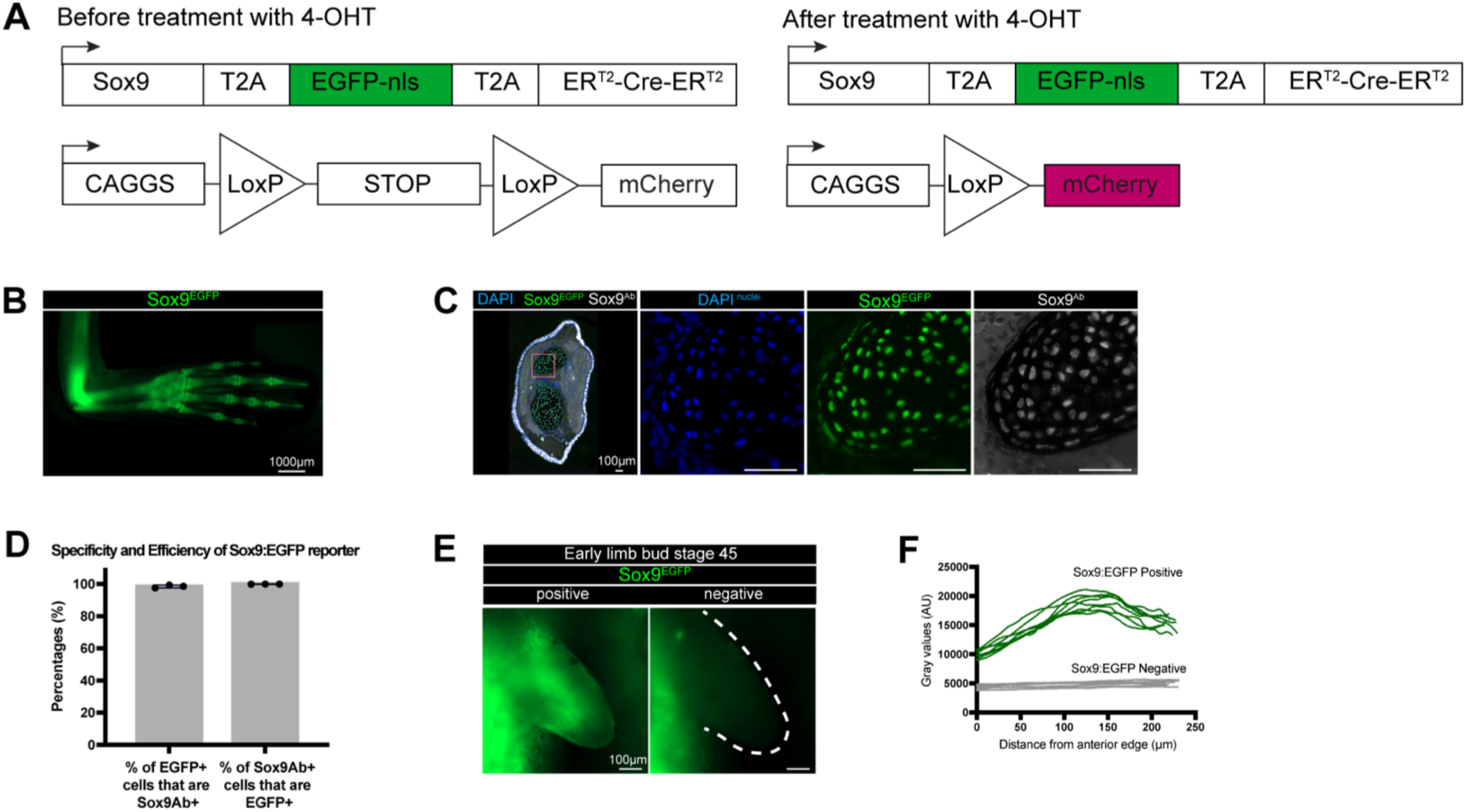
Characterization of *Sox9*:EGFP-nls-T2A-ER-Cre-ER transgenic reporter. **A** Schematic of *Sox9* transgenic reporter design and fate mapping strategy. Upon 4-OHT induction, the STOP cassette is removed and *Sox9* positive cells and their progeny express mCherry. **B** Widefield image of a limb from the *Sox9*:EGFP reporter (green) (representative of *n* = 4 animals, 2 limbs each). **C** Cross-section of the lower limb of a *Sox9* transgenic stained with DAPI (nuclei, blue) and anti-SOX9 antibody (white). EGFP (green) signal reflects endogenous fluorescence. Merged image is on the left. The boxed region in the left panel is magnified to the right. **D** Quantification of the specificity and efficiency of the *Sox9*:EGFP reporter using cross-sections in (C). Each dot represents data from one limb. Specificity: the percentage of EGFP positive cells in the skeleton that stained with SOX9 antibody, normalized to the number of sections per limb. Efficiency: the percentage of SOX9 antibody positive cells in skeleton that were positive for the *Sox9*:EGFP reporter, normalized to the number of sections per limb (*n* = 3 limbs). **E** Widefield image of a stage 45 limb bud from the *Sox9*:EGFP (green) transgenic. Left: positive animal; Right: negative control. **F** Quantification of EGFP (green) signal gray values along the anterior-to-posterior axis, quantified from images in (E) (*n* = 8 animals, 1 limb bud each).

Live imaging revealed the expected expression of EGFP in cartilage skeleton at steady state (Figure 1B) and during regeneration (Figure S1A). To confirm that EGFP reflected *Sox9* expression, we harvested uninjured limbs from *Sox9* transgenic axolotls, prepared cross- sections and stained these with anti-SOX9 antibody (Figure 1C). 98.5% ± 0.9 of EGFP- expressing cells in the skeleton stained with SOX9 antibody, demonstrating high specificity of the reporter, while 100% of antibody-stained cells in the skeleton were EGFP positive indicating high efficiency (Mean *n* = 344 cells per section, from 8 sections, from 3 limbs) (Figure 1D). In addition to expression in the skeleton, we observed EGFP in epidermal cells. Analyzing published single cell RNA-seq datasets revealed that both mesenchymal cells and epidermal cells express *Sox9* in axolotls and in mice (McMann et al., 2025) (Figure S1B). Thus, our *Sox9* transgenic line faithfully reports *Sox9* expression in the limb.

To profile *Sox9* expression during limb development, we imaged animals at an early limb bud stage (stage 45) (Figure 1E) (*n* = 8 animals, 1 limb bud each), a mid limb bud stage (stage 47) (Figure S1C) (*n* = 4 animals, 1 limb bud each) or at 4 cm size after completing limb development (Figure 1B) (*n* = 4 animals, 2 limbs each). We found that EGFP in early limb buds is expressed most strongly in the center of the bud, in the condensing cartilage region, but that it is expressed at lower levels more broadly throughout the mesenchyme and epidermis (Figures 1E-F) (*n* = 8 animals, 1 limb bud each). Such wide expression was not evident in fully developed limbs (Figure 1C, *n* = 8 sections, from 3 limbs). These results suggest that although *Sox9* expression is initially broader, it becomes restricted to skeletal elements as limb development proceeds.

### Progressive restriction of *Sox9* expression in the developing axolotl limb

To define the contributions of *Sox9*-expressing cells to limb development, we performed genetic lineage tracing. Crossing the *Sox9* transgenic line with our existing memory cassette axolotl (Kawaguchi et al., 2024) allowed us to permanently convert *Sox9*-expressing cells with mCherry by treating with the tamoxifen analogue 4-hydroxytamoxifen (4-OHT) at the desired time (Figure 1A). We induced conversion in the early limb bud (stage 45) (Figure S2A), mid limb bud (stage 47/48) (Figure S2A) or at both early and mid-limb bud stage (Figure 2A), then raised the animals up to 4 cm size to complete limb development (‘limb bud converted’ cohorts). As a comparison, we induced conversion directly in the fully developed limb of 4 cm animals (‘mature converted’ cohort) and waited 7 days before imaging (Figure 2A). Although epidermis was converted in these experiments, we focus our descriptions on mesenchymal cells. It was previously shown that epidermis is a separate lineage compartment that does not contribute to mesenchyme or skeleton (Kragl et al., 2009).

**Figure 2.**
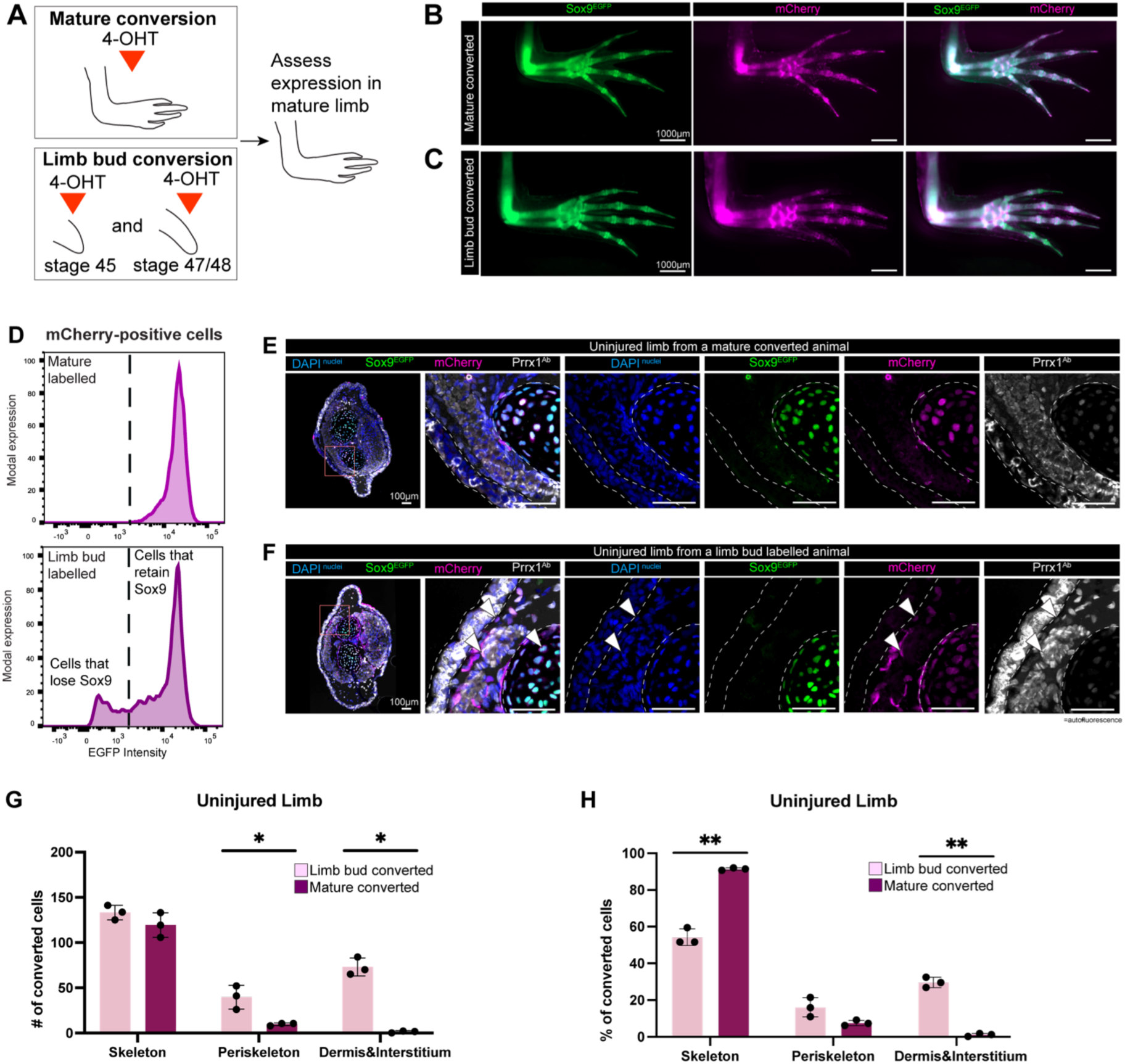
Progressive restriction of *Sox9* expression in the developing axolotl limb. **A** Schematics indicating the timing of conversion in lineage tracing experiments. After treatment with 4- OHT, animals were raised to 4 cm. **B** Widefield images of mature converted limbs (*n* = 9 animals, 2 limbs each). **C** Widefield images of limb bud converted limbs (*n* = 7 animals, 2 limbs each). **D** Flow cytometry analysis of mature converted limbs (upper panel) or limb bud converted limbs (lower panel). Panels depict EGFP fluorescence intensity in mCherry-positive cells (*n* = 3 animals, 2 limbs each, per condition). **E** Cross-sections of an uninjured limb from a mature converted animal. Images show DAPI (nuclei, blue), *Sox9*:EGFP reporter (green), mCherry conversion (magenta) and PRRX1 antibody staining (white). **F** Cross-sections of an uninjured limb from a limb bud converted animal. Arrowheads indicate converted cells in dermis and interstitium compartments. Images show DAPI (nuclei, blue), *Sox9*:EGFP reporter (green), mCherry conversion (magenta) and PRRX1 antibody staining (white). **G** Quantification of mCherry positive cells in different compartments (skeleton, periskeleton, dermis and interstitium), normalized to the number of sections quantified per limb. Each dot represents data from one limb (*n* = 3 limbs per condition). Bars show mean ± SD. Differences between conversion time points were assessed for each compartment (two-tailed unpaired Welch’s t-test on log₁₀(Y+1)-transformed values with Holm–Šídák correction for multiple comparisons; skeleton: adjusted P = 0.228281, t = 1.506, df = 3.033; periskeleton: adjusted P = 0.024011, t = 6.050, df = 2.708; dermis/ interstitium: adjusted P = 0.017270, t = 10.7, df = 2.226; *n* = 3 limbs per condition. **H** Percentage of mCherry positive cells in different compartments (skeleton, periskeleton and dermis and interstitium) normalized to the number of sections quantified per limb. Each dot represents data from one limb (*n* = 3 limbs per condition). Bars show mean ± SD. Differences between time of conversion were assessed for each compartment (two-tailed unpaired Welch’s t-test on arcsine square-root transformed percentage with Holm–Šídák correction for multiple comparisons; skeleton: adjusted P = 0.004315, t = 16.39, df = 2.256; periskeleton: adjusted P = 0.071072, t = 2.974, df = 2.580; dermis/ interstitium: adjusted P = 0.001091, t = 14.44, df = 3.393; *n* = 3 limbs per condition.

We first described the overall pattern of conversion using low resolution widefield microscopy. As expected, mature converted limbs displayed high apparent overlap between EGFP signal, permanent mCherry conversion and skeleton in all forelimb segments (upper arm, lower arm and hand, including digits) (Figure 2B). Next, we analyzed limb bud converted animals. Early converted limbs (Figure S2B), mid converted limbs (Figure S2C) and double converted limbs (Figure 2C) also displayed apparent overlap between EGFP, permanent mCherry conversion and skeleton, although the extent of mCherry conversion in the skeleton differed depending on the timing of treatment with 4-OHT. *Sox9* cells that were converted in the early limb bud ultimately contributed to upper arm, lower arm and hand plate skeleton, while those converted at mid limb bud stages additionally contributed to the anterior digits (Figures S2B-C). Double converted limbs displayed a cumulative signal, with more complete conversion and additional signal in posterior digits (Figure 2C). These results align with the order of skeleton differentiation in the limb: proximal segments (upper arm and lower arm) first, followed by more distal segments (hand plate, anterior digits then posterior digits) (Riquelme-Guzmán et al., 2022a).

Sibling animals that did not receive 4-OHT treatment showed no conversion, confirming the specificity of mCherry induction (Figure S2D).

### A minority of cells loses *Sox9* expression during limb development

We used the converted animals to estimate when, and how many, cells lose *Sox9* expression during limb development (Figures 1C and 1E). We harvested the lower arm and hand segments of fate mapped limbs, dissociated these into single cells and analyzed EGFP and mCherry expression using flow cytometry. EGFP acts as a readout for current *Sox9* expression, while mCherry labels cells that expressed *Sox9* at the time of 4-OHT treatment (Figure 1A). We expected to detect *Sox9*-expressing cells that later switched off *Sox9* as mCherry-positive cells that are not EGFP-positive in the fully developed limb.

As a reference, we first analyzed mature converted limbs. We found that 99.8% of mCherry- positive cells were EGFP-positive (Figure 2D, *n =* 9863 mCherry cells pooled from 3 animals, 2 limbs each), consistent with stable EGFP expression. The conversion efficiency in this experiment (percentage of EGFP-positive cells converted with mCherry) was 52.4%.

Next, we analysed limb bud converted limbs. Interestingly, in limbs double converted during development, 19.4% of mCherry-positive cells were negative for EGFP (Figure 2D, *n =* 9834 mCherry cells pooled from 3 animals, 2 limbs each). This shows that some cells that previously expressed *Sox9* in the limb bud subsequently shut down expression. We found that 24.2% of mCherry-positive cells were EGFP-negative in early limb bud converted animals (Figure S2E, *n =* 9839 mCherry cells pooled from 2 animals, 2 limbs each), while this percentage dropped to 10.8% in mid limb bud converted animals (Figure S2E, *n =* 9899 mCherry cells pooled from 3 animals, 2 limbs each) supporting a progressive shutdown process, Together, these results support that a minority of cells expressing *Sox9* during limb development shuts down expression before development is complete.

### Cells that lose *Sox9* expression during development become interstitial and dermal fibroblasts

Next, we sought to identify the *Sox9* lineage cells in mature converted samples and limb bud converted samples (early + mid double conversion). We fixed converted limbs and analyzed tissue cross-sections, quantifying only the mesenchymal compartment. We found the extent of mCherry conversion between biological replicates to be consistent, with more conversion found in limb bud converted samples that were twice treated with 4-OHT (Figure S2F). The mean number of mCherry-positive cells per tissue section in mature-converted limbs was 131 ± 14 (*n* = 13 tissue sections from 3 limbs), compared to 246 ± 11 converted cells per tissue section in limb bud converted limbs (*n* = 6 tissue sections from 3 limbs) (Figure S2F).

We categorized mCherry-positive cells into ‘skeleton’ (cartilage), ‘periskeleton’ (perichondrial cells surrounding the cartilage) or ‘dermis and interstitium’ (fibroblasts residing in the skin or muscle compartments respectively). In mature converted samples, mCherry-positive cells were almost exclusively skeletal or periskeletal, with just 1% found in the dermis or interstitium (Figures 2E, 2G-H). By contrast, limb bud converted samples had a significantly higher percentage of mCherry-positive cells in dermis or interstitium, at 30% (Figures 2F-H). These cells were fibroblasts, as confirmed by their morphology, location and antibody staining against the homeodomain transcription factor PRRX1 (Figure 2F) (Gerber et al., 2018).

Together, our flow cytometry and tissue section-based analyses reveal that not all cells that express *Sox9* during development become skeleton or skeleton-associated. Instead, some cells lose *Sox9* expression and become loose mesenchymal cells including dermal and interstitial fibroblasts.

### Cells that retain or lose *Sox9* expression during development contribute differently to limb regeneration

Next, we queried whether *Sox9* lineage cells that retain or lose *Sox9* expression during development contribute differently to regeneration. Previous studies suggest that skeletal cells contribute only minimally to regeneration, by forming a callus, while dermal and interstitial fibroblasts can regenerate new skeleton (Gerber et al., 2018). However, these lineage tracing experiments were not performed specifically in the context of the *Sox9* lineage.

To induce regeneration, we performed an amputation through the top third of the lower arm of fate mapped animals. We performed live imaging of regenerating limbs by widefield microscopy at 6, 16 and 43 days post amputation (dpa). These time points reflect early blastema (6 dpa), mid-regeneration when regenerating skeleton is visible (16 dpa) and after full regeneration (43 dpa). Although we report the fate mapping results in the limb mesenchyme, we also observed patches of mCherry-converted cells in the epidermis, presumably arising from epidermal cells converted before the injury.

We found that *Sox9*-expressing cells in the mature limb hardly contribute new cells to regeneration. mCherry cells were predominantly restricted to the tip of the amputated skeleton at 6 dpa (*n* = 9 animals, 2 limbs each), 16 dpa (*n* = 6 animals, 2 limbs each) and in the fully regenerated limb (*n* = 6 animals, 2 limbs each) (Figure 3A). Comparing mCherry signal with the *Sox9*:EGFP reporter reinforced the relative absence of converted cells in the regenerated skeleton. By contrast, amputation of limb bud converted limbs resulted in a widespread contribution of *Sox9* lineage cells to the regenerated skeleton and to other mesenchymal compartments at all time points (*n* = 4 animals, 2 limbs each) (Figure 3B).

**Figure 3.**
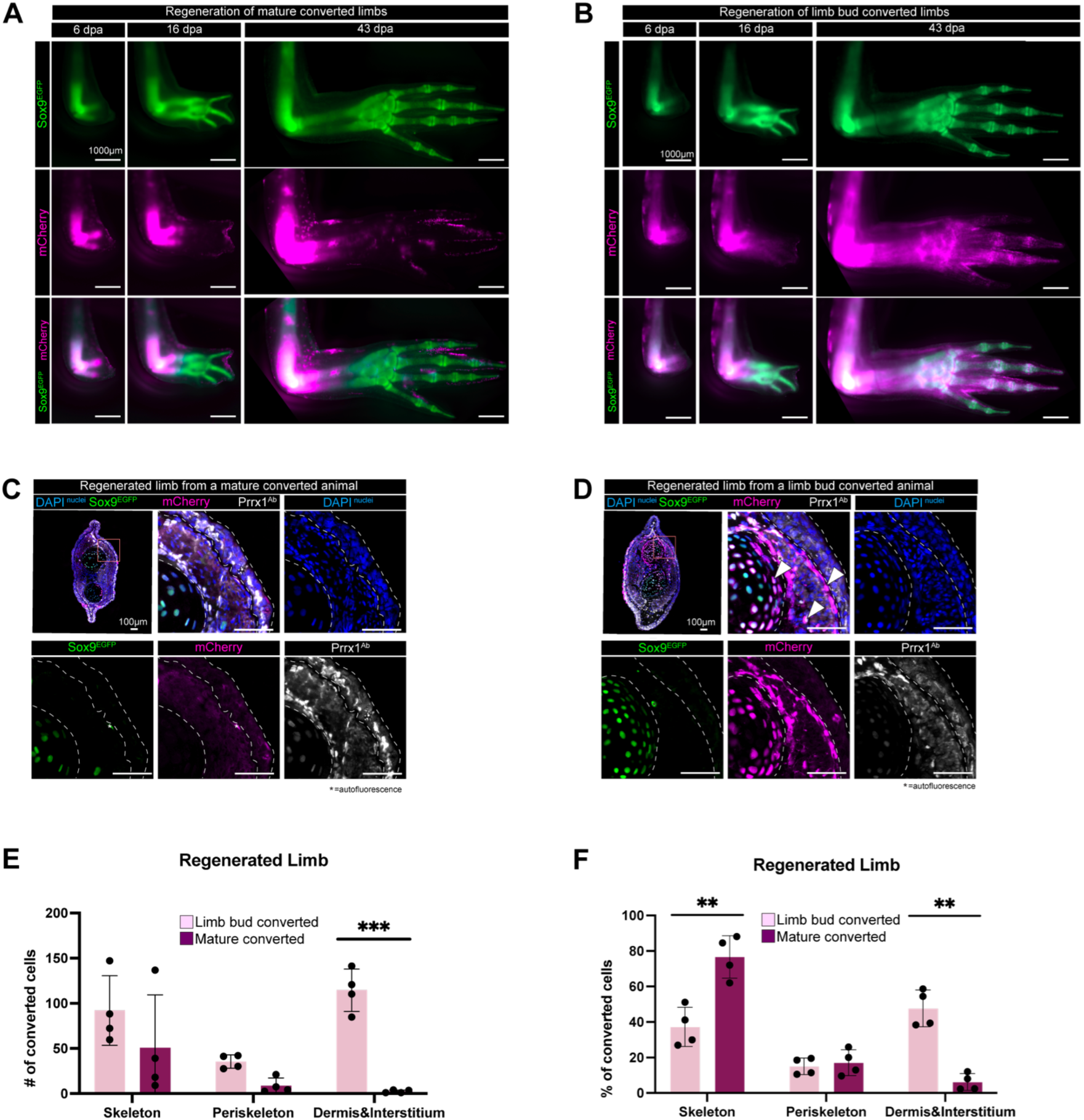
Cells that shut down *Sox9* expression contribute extensively to limb regeneration. **A** Widefield images of a mature converted limb undergoing lower arm regeneration (Representative of *n* = 6 animals, 2 limbs each). **B** Widefield images of a limb bud converted limb undergoing lower arm regeneration (Representative of *n* = 4 animals, 2 limbs each). **C** Cross-section of a regenerated limb from the mature converted cohort (A). Images show DAPI (nuclei, blue), *Sox9*:EGFP reporter (green), mCherry conversion (magenta) and PRRX1 antibody staining (white). **D** Cross-section of a regenerated limb from the limb bud converted cohort (B). Arrowheads indicate converted mCherry cells in dermis, interstitium and skeleton. Images show DAPI (nuclei, blue), *Sox9*:EGFP reporter (green), mCherry conversion (magenta) and PRRX1 antibody staining (white). **E** Number of mCherry positive cells contributing to different compartments (skeleton, periskeleton, dermis and interstitium) of the regenerated limb when converted in the mature limb or in the limb bud. Each dot represents one limb, normalized to the number of sections analysed (2–6 per limb); Bars show mean ± SD. Differences between conversion conditions were assessed for each compartment (two-tailed unpaired Welch’s t-test on log₁₀(Y+1)-transformed values with Holm–Šídák correction for multiple comparisons; skeleton: adjusted P = 0.171584, t = 1.694, df = 3.687; periskeleton: adjusted P = 0.051793, t = 3.764, df = 3.409; dermis/ interstitium: adjusted P = 0.000522, t = 13.26, df = 4.047; *n* = 4 limbs per condition.) **F** Percentage of mCherry positive cells in different compartments (skeleton, periskeleton, dermis and interstitium) of the regenerated limb when converted in the mature limb or in the limb bud, normalized to the number of sections per limb. Each dot represents data from one limb. Bars show mean ± SD. Differences between time of conversion were assessed for each compartment (two-tailed unpaired Welch’s t-test on arcsine square-root transformed percentage with Holm–Šídák correction for multiple comparisons; skeleton: adjusted P = 0.008152, t = 4.616, df = 5.729; periskeleton: adjusted P = 0.682970, t = 0.4318, df = 5.282; dermis/interstitium: adjusted P = 0.001046, t = 7.264, df = 5.990; *n* = 4 limbs per condition)

To evaluate the spatial distribution and contributions of mCherry converted cells at the cellular level, we harvested fully regenerated limbs at 43 dpa and prepared tissue cross-sections from the regenerated part for further analysis (Figures 3C-D). The mean number of mCherry-positive cells per section in mature-converted regenerates was 62 ± 68 (*n* = 23 sections, from 6 limbs), compared to 241 ± 33 converted cells per section in regenerates harvested from limb bud converted limbs (Figure S3A) (*n* = 16 sections, from 4 limbs). Of the cells regenerated by mature converted *Sox9* cells, 94% ± 5 were skeletal or periskeletal cells and 6% ± 5 were dermal or interstitial fibroblasts (Figures 3E-F). The corresponding percentages for limb bud converted *Sox9* cells were 52% ± 10 skeletal or periskeletal cells and 48% ± 10 dermal or interstitial fibroblasts (Figures 3E-F).

Our analyses demonstrate divergent behaviours within the *Sox9* lineage. Cells retaining *Sox9* expression post-development contribute in a limited way to limb regeneration, primarily regenerating skeleton and peri-skeleton. Meanwhile, cells that lose *Sox9* expression contribute extensively to skeletal regeneration and additionally regenerate dermal and interstitium fibroblasts that reside further away from the skeleton.

### Transiently *Sox9*-expressing cells are also produced during limb regeneration

Our experiments have shown that *Sox9* positive cells in the limb bud generate different cell populations during limb development. We queried whether a similar mechanism is deployed during regeneration. To test this possibility, we prepared *Sox9* lineage tracing animals, raised these to 5 cm, performed limb amputation through the lower arm then treated with 4-OHT to induce conversion specifically during regeneration. We treated with 4-OHT at 6 dpa (early regeneration) or 26 dpa (late regeneration, when skeleton has re-formed) (Figure 4A, S4A), then waited until 33 dpa to allow sufficient time for Cre/loxP recombination and expression of mCherry. Both treatments resulted in converted cells in the fully regenerated limb (Figure 4B-C), analogous to the results of our limb bud lineage tracing experiments.

**Figure 4.**
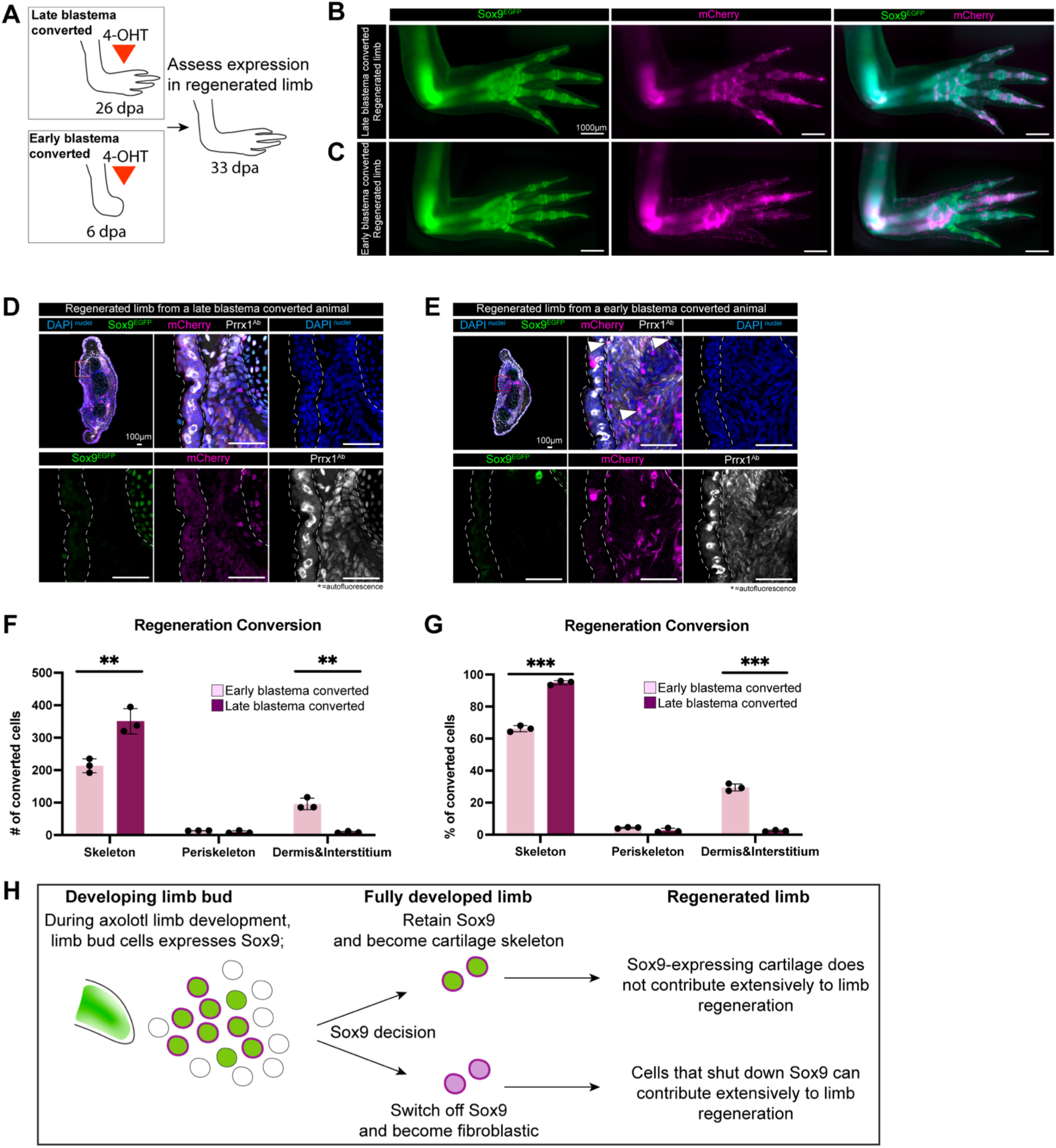
Transiently *Sox9*-expressing cells are also produced during early regeneration. **A** Schematics indicating the timing of conversion with 4-OHT: at late blastema stage (26 dpa, upper panel) or at early blastema stage (6 dpa, lower panel). After treatment, animals were left to regenerate their limbs up to 33 dpa. **B** Widefield images of a regenerated limb converted at late blastema stage (Representative of *n* = 5 animals, 2 limbs each). **C** Widefield images of a regenerated limb converted at early blastema stage (Representative of *n* = 4 animals, 2 limbs each). **D** Cross-sections of a regenerated limb (33dpa) from a late blastema converted animal. Images show DAPI (nuclei, blue), *Sox9*:EGFP reporter (green), mCherry conversion (magenta) and PRRX1 antibody staining (white). **E** Cross-sections of a regenerated limb (33dpa) from an early blastema converted animal. Images show DAPI (nuclei, blue), *Sox9*:EGFP reporter (green), mCherry conversion (magenta) and PRRX1 antibody staining (white). Arrowheads indicate converted mCherry cells in skeleton, dermis and interstitium compartments. Images show DAPI (nuclei, blue), *Sox9*:EGFP reporter (green), mCherry conversion (magenta) and PRRX1 antibody staining (white). **F** Quantification of the contribution of mCherry positive cells from early or late blastema converted animals to different compartments (skeleton, periskeleton, dermis and interstitium), normalized to the number of sections quantified per limb. Each dot represents data from one limb (*n* = 3 limbs per condition). Bars show mean ± SD. Differences between conversion time points were assessed for each compartment (two-tailed unpaired Welch’s t-test on log₁₀(Y+1)-transformed values with Holm–Šídák correction for multiple comparisons; skeleton: adjusted P = 0.008957, t = 5.799, df = 3.973; periskeleton: adjusted P = 0.209779, t = 1.816, df = 2.021; dermis/interstitium: adjusted P = 0.001239, t = 12.81, df = 3.568; *n* = 3 limbs per condition). **G** Percentage of contribution of mCherry positive cells from early or late blastema converted animals to different compartments (skeleton, periskeleton, dermis and interstitium) normalized to the number of sections quantified per limb. Each dot represents data from one limb (*n* = 3 limbs per condition). Bars show mean ± SD. Differences between time of conversion were assessed for each compartment (two- tailed unpaired Welch’s t-test on arcsine square-root transformed percentage with Holm–Šídák correction for multiple comparisons; skeleton: adjusted P = 0.000203, t = 19.42, df = 3.592; periskeleton: adjusted P = 0.166916, t = 2.016, df = 2.252; dermis/ interstitium: adjusted P = 0.000203, t = 25.27, df = 3.303; *n* = 3 limbs per condition). **H** Model linking *Sox9* expression dynamics during limb development with contributions to limb regeneration.

If a similar mechanism is used as during development, we might expect that *Sox9* cells converted during early regeneration give rise to skeleton, periskeleton, dermis and interstitium, while those converted later during regeneration only give rise to skeleton and periskeleton. We found this to be the case. We quantified serial tissue sections harvested from regenerated limbs at 33 dpa. *Sox9*-expressing cells converted in the late blastema contributed almost exclusively to skeleton or periskeleton (98% ± 0.5 of cells) (Figures 4D, 4F and 4G, mean *n* = 369 converted cells per section, 10 sections, from 3 limbs). Meanwhile, when *Sox9*-expressing cells were converted in the early blastema, 71% ± 2.2 of the resulting cells were skeletal or periskeletal, while 29% ± 2.2 were dermal or interstitium (Figures 4E-G, mean *n* = 323 converted cells per section, 8 sections, from 3 limbs). The number of converted cells counted per limb was similar in both experimental groups (Figure S4B).

In sum, our fate mapping experiments suggest that transiently *Sox9*-expressing cells contribute both to limb development and to limb regeneration.

## Discussion

Genetic fate mapping allows tracing of molecularly defined cell populations and their progeny over time, revealing their contributions to developing tissues. Although SOX9 transcription factor is classically associated with cartilage formation in mesenchymal tissues, fate mapping revealed wider contributions to the skeletal and periskeletal components of cartilage and bone, as well as ligaments, tendons and synovia in mouse limb buds (Akiyama et al., 2005; Soeda et al., 2010). Timed conversion revealed stage-specific behaviours: for example, *Sox9* lineage cells separate into osteoblast- and chondrocyte-forming lineages prior to E12.5 (Soeda et al., 2010) and *Sox9*- expressing cells converted at E10.5 produced more perichondrial cells than those converted at either E8.5 or E12.5 (Wu et al., 2025). More cell types are converted after *Sox9* lineage tracing than by a reporter reflecting direct *Sox9* expression at E17.0 (Akiyama et al., 2005). This suggests that some cells that initially express *Sox9* during limb development lose *Sox9* expression and contribute to non-cartilage tissues. However, whether such a strategy is used during limb development outside of the mouse was not known, and the full range of cell types that derive from initially *Sox9*-expressing progenitors remains incompletely understood.

Axolotls offer an opportunity to examine *Sox9* lineage decisions during limb development and during limb regeneration. Although *Sox9* expression has been visualized in axolotls using *in situ* hybridisation, antibody staining and genetic reporters, fate mapping had not yet been performed (Purushothaman et al., 2021; Riquelme-Guzmán et al., 2022b). Here, we fate mapped *Sox9*- expressing cells during axolotl limb development and regeneration. We show that cells that express *Sox9* during early limb development or regeneration make a choice: to retain stable *Sox9* expression and become cartilage or perichondrial cells or to lose *Sox9* expression and contribute to other cell types such as dermal or interstitial fibroblasts (Figure 4G). When we converted *Sox9*-expressing cells in the early limb bud and traced their progeny into the fully developed limb, we found that almost 25% no longer expressed *Sox9*. Our time-resolved experiments suggest that cells gradually shut off *Sox9* expression throughout limb bud development rather than at a defined developmental stage in a synchronized fashion.

Previously, it was shown that there is an on-to-off dynamic of *Sox9* expression in mouse limb buds, with previously *Sox9*-expressing cells contributing not only to skeleton but also to skeleton-associated mesenchyme (Akiyama et al., 2005). In axolotls, we observed that these cells can also contribute to the dermal compartment. Given how far away these cells are from the limb skeleton, this demonstrates high plasticity in this cell population. Whether this is an axolotl-specific feature or was not reported in the mouse fate mapping experiments is not clear.

We show that cells that retain or lose *Sox9* expression contribute differently to limb regeneration. Cells that retain *Sox9* become cartilage and peri-skeleton that regenerate only a limited amount of tissue after injury. By contrast, cells that lose *Sox9* expression contribute significantly to mesenchymal tissues in the regenerated limb, including re-initiating *Sox9* expression and forming new skeleton (Figure 4G). These results are consistent with the prevailing model that *Prrx1*-expressing fibroblasts, and not skeletal cells, are a multipotent source of cartilage, bone, tendons, ligaments and fibroblasts during limb regeneration (Gerber et al., 2018). An intriguing possibility is that previous expression of *Sox9* acts as a cellular ‘memory’ to promote re-expression and cartilage formation during regeneration, even if *Sox9* expression was shut down in the interim. Our experiments do not distinguish if cells can lose and gain *Sox9* expression multiple times during development - for example in an oscillatory manner as can occur in synchronized limb micromass cultures *in vitro* (Alagha et al., 2021; Vágó et al., 2023). Nevertheless, cells can repeat the ‘*Sox9* decision’ during regeneration (Figure 4) - potentially resulting in many cycles throughout an axolotl’s lifetime.

As part of our characterizations of the *Sox9* transgenic line, we observed *Sox9* lineage cells in the epidermis. Converted cells in the epidermis appeared to generate new epidermal cells during regeneration, as previously reported (Kragl et al., 2009). Interestingly, epidermal *Sox9* cells were not described in a previous study that generated a *Sox9*:mCherry knock-in axolotl line (Riquelme-Guzmán et al., 2022b), nor in *in situ* hybridisations targeting *Sox9* transcripts during axolotl limb bud development (Purushothaman et al., 2021). Nevertheless, the presence of *Sox9*-expressing cells in epidermis is evident in gene expression datasets of axolotl limb development (McMann et al., 2025). In mice, *Sox9*-expressing cells can contribute to several skin-associated compartments such as sweat glands and ducts in the paw pads (Lu et al., 2012), hair follicles and sebaceous glands in the tail skin (Nowak et al., 2008) and to the nail bed epithelium (Lao et al., 2022). Thus, a function for *Sox9* in the skin compartment may be observed in both axolotls and mice, but this may be evolutionarily divergent given that axolotls do not possess sweat glands, hair follicles or nails.

It will be important to delineate the molecular and spatiotemporal regulation of *Sox9* lineage cells. Lineage tracing in the developing mouse limb has revealed that distinct *Sox9*-expressing sub-populations generate different progeny. For example, bone eminences, specialised structures where tendon-to-bone attachment take place, originate from a secondary pool of progenitors that are *Sox9*- and *Scx*-positive but *Col2a1*-negative at E13.5 (Blitz et al., 2013).

Recent work has started to unravel the molecular regulation of *Sox9*-expressing sub- populations. CUT&RUN revealed differences in the genomic binding of SOX9 in different mouse limb bud progenitors across time (Sega et al., 2026). It will be interesting to compare the molecular regulation of diverse *Sox9* lineage cells in mouse and axolotl, particularly given the high degree of plasticity in the axolotl, in which previously *Sox9*-expressing cells can become *Sox9*-negative dermal fibroblasts residing under the skin.

In sum, fate mapping of *Sox9*-expressing cells during axolotl limb development and regeneration revealed that there is a transiently *Sox9*-expressing population in the early limb bud and blastema that subsequently shuts down *Sox9* and gives rise to non-skeleton- associated cells including dermal fibroblasts. Cells that lose *Sox9* expression show greater contributions to limb regeneration than cells that retained *Sox9* expression. These results link the expression dynamics of *Sox9* to different cell fates and outcomes during regeneration.

## Materials and methods

### Ethics oversight

All the animal experiments were conducted according to ethical guidelines and approved by the Magistrate of Vienna (Genetically Modified Organism Office and MA58, City of Vienna, Austria), under licences GZ: 51072/2019/16, GZ: MA58-1432587-2022-12 and GZ: MA58-1516101- 2023-21, 2024-0.438.721 and 2024-0.438.718.

### Axolotl (*Ambystoma mexicanum*) husbandry

Axolotl breedings and maintenance were performed by the animal caretaker team at IMP/IMBA animal facilities. Animals were raised in individual tanks in Vienna tap water. Live imaging and limb amputations were performed under anesthesia in 0.015% benzocaine (Merck, E-1501) diluted in Vienna tap water (Khattak et al., 2014). After limb amputation, animals were treated with butorphanol analgesic solution (0.5mg/L) for 72 hours. At experimental endpoint, animals were euthanized by anaesthetic overdose.

We performed each experiment with sibling animals from the same clutch, randomly assigned to different experimental conditions and reared under matching conditions throughout the experiments. Axolotl sizes were measured from snout to tail. Axolotls up to 10 cm in length were used in experiments. Limb amputation was performed on forelimbs through the upper-third of the zeugopod (lower arm), unless indicated otherwise.

### Genetic fate mapping

We used the *Sox9*:EGFP/lineage tracing line reported by Tardivo and Olmeda et al., (2026): *tm(Sox9t/+:Sox9-eGFPnls-T2A-ERT2-Cre-ERT2)Etnka*.

To perform *Sox9* lineage tracing, we crossed this line with our previously published *Caggs:loxP- GFP-dead(Stop)-loxP-mCherry)Etnka* memory cassette line (Kawaguchi et al., 2024). We fluorescently screened progeny animals for *Sox9*:EGFP expression using an AXIOzoom V16 widefield microscope (Zeiss) and confirmed that they were negative for mCherry expression, which would reflect spurious Cre-loxP recombination.

To induce conversion by Cre-loxP recombination, we bathed animals in 100 mL of 1 µM 4-OHT (for developing animals) or 2 µM 4-OHT (for mature or regenerating animals) for 16 hours in the dark according to the method of Khattak et al., (2014). For limb bud conversion, we treated with 4-OHT at stage 45 (early) or stage 46/47 (mid) or at both stages (double conversion, with animals returned to water between 4-OHT treatments). Subsequently, these animals were raised in individual tanks up to 4 cm size. We performed mature conversion on sibling animals from the same clutch as the limb bud conversion experiments, raised to 4cm. For regeneration conversion, we used a separate clutch of animals raised to 5 cm.

All animals were screened under anaesthesia at least 7 days after 4-OHT treatment to allow time for Cre-loxP recombination and mCherry expression using an AXIOzoom V16 widefield microscope (Zeiss).

### Flow cytometry

Fate mapped limbs were harvested distally from the original amputation plane including the hand (no digits) and dissociated in Liberase TM enzyme (Merck, 05401127001). The dissociation protocol was adapted from the previously published method (Lin et al., 2021), with the following changes: Dissociation solution was prepared in a serum-free axolotl cell culture media. Cell culture media was adopted from previously published study (Chen et al., 2011): 60% Advanced DMEM/F12 (Thermo Fisher Scientific, 12634010) diluted based media supplemented with 1% GlutaMAX™ Supplement (Thermo Fisher Scientific, 35050061), 1% Insulin-Transferrin-Selenium (ITS-G) (Thermo Fisher Scientific, 41400045), 1% L-Ascorbic acid 2-phosphate sesquimagnesium salt hydrate (Sigma Aldrich, A8960), 1% MEM Non-Essential Amino Acids Solution (Thermo Fisher Scientific, 11140035), 1% Penicillin-Streptomycin (Thermo Fisher Scientific, 15140122). After cutting the tissue into small pieces, dissociation was performed for 4 hours at 25°C degrees. Subsequently, cells were filtered through a 70 μm cell strainer (Miltenyi Biotech, 130-098-462). Cells were run through a flow cytometer (BD Fortessa) using a 100μm nozzle. Quantification and analyses were performed using FlowJo software (BD Biosciences).

### Cryosectioning, immunostaining and imaging

Limb samples were harvested and fixed overnight in 4% formaldehyde/PBS at 4°C followed by thorough washing of samples with cold PBS. Samples were prepared for cryosectioning by incubating in 20% sucrose in PBS overnight followed by 30% sucrose in PBS overnight at 4°C. Samples were transferred into 100% Tissue-Tek O.C.T compound and acclimatized to room temperature at least for 1 hour before embedding. Samples were mounted in fresh Tissue-Tek O.C.T. compound, frozen on dry ice then stored at -70°C. Cryosectioning was performed using a Cryostar NX70 (Thermo Fisher Scientific), with sections collected at 16 µm thickness on Permaflex adhesion slides (Leica, 3800002). Slides were stored at -20°C or -70°C for short or long-term storage, respectively.

For immunostaining, slides were acclimatized to room temperature, in the dark, for 30 minutes to 1 hour. All following steps were performed at room temperature unless specified otherwise. Slides were washed with PBTx (0.2% TritonX-100 in PBS) thoroughly before starting the staining. For DAPI-only staining: washed slides were incubated for 30 mins with 5 µg/mL DAPI in PBTx followed by PBTx washes before mounting. For antibody staining: washed slides were blocked with blocking solution (3% BSA in PBTx) for 30 mins. After blocking, slides were incubated with primary antibody solution overnight at 4°C. The following primary antibodies were used: rabbit anti-PRRX1 at 1:200 (Gerber et al. 2018); rabbit anti-SOX9 at 1:500 (R&D Systems, AF3075) diluted in 0.3% BSA in PBTx. Slides were washed thoroughly with PBTx and incubated with secondary antibody solutions overnight at 4°C: anti-rabbit Alexa-647 (Invitrogen, A-21244) 1:500 diluted in PBTx. Slides were washed thoroughly again with PBTx. Slides were mounted in AbberiorMount liquid antifade mounting media. Slides were imaged using a Pannoramic 250 FLASH III Digital Scanner.

### Image analysis

Images were analyzed using CaseViewer version2.4 (3dhistech, RRID: SCR_017654), or Fiji software version 2.16.0 (Schindelin et al., 2012).

### Quantification and statistical analyses

Converted cells were manually quantified in 2 – 6 sections per limb using the CaseViewer version2.4 (3dhistech, RRID: SCR_017654) counting tool . Quantified cells were summed up across different sections of the same limb. To control for variations in the number of sections analyzed per limb, raw counts were normalized to number of sections and represented as number of cells per section. Percentages were calculated by normalizing the total number of converted cells in a compartment (skeleton, periskeleton or dermis and interstitium) to the total number of converted cells across all compartments. Mean values are given ± standard deviation (SD).

Statistical analyses were carried out using Prism version 11.0.2 (GraphPad). Data were evaluated for normality in distribution prior to statistical analysis. Proportional data were arcsine square-root transformed (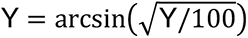) before analysis. Absolute compartmental counts were transformed using Y = log(Y + 1) to account for zero values, and total cell counts per limb were log-transformed (Y = log(Y)). Grouped comparisons of proportions and compartmental counts between treatments were performed using unpaired Welch’s *t*-tests (not assuming equal standard deviations) followed by Holm–Šídák multiple comparisons correction. Total cell counts per section were compared using an unpaired Welch’s *t*-test. Figures display untransformed percentages or normalized cell counts. Significance thresholds were set at *α* = 0.05, with adjusted *p*-values denoted as: ns, not significant (*p* ≥ 0.05); *, *p* < 0.05; **, *p* < 0.01; ***, *p* < 0.001; ****, *p* < 0.0001. Statistical details, including exact sample numbers (*n*), are specified in the respective figure legends.

### Single-cell RNA sequencing analyses

We used data from a single cell RNA sequencing atlas from mouse and axolotl hindlimbs (McMann et al., 2025). An interactive version of processed scRNA-seq datasets was available at *digilimb.wi.mit.edu*. We searched for the *Sox9* gene in both axolotl and mouse datasets and reported the *Sox9* expression levels in the corresponding UMAP for each species.

## Acknowledgments

We thank A. Fischer for establishing changes to the tissue dissociation protocol and the animal care takers team (A. Lentz-Koblenc, D. Bastian, J. König, M. Sorgner, V. Szilagyi) for axolotl care. We thank staff at the Histology facility (A. Piszczek, M. Grivej, T. Engelmaier) and staff at the BioOptics facility (A. Moreno Cencerrado, G. Schmauss, G. Bradamante, J. Sonntag, M. Weninger, P. Pasierbek) for their support.

## Open access

This research was funded in part by European Research Council (ERC) Advanced Grant 742046. For the purpose of open access, the authors have applied a CC BY public copyright licence to any Author Accepted Manuscript version arising from this submission.

## Author contributions

M.N.B designed experiments, performed all experiments, analyzed data and wrote the manuscript. L.O conceived the project, secured funding, designed experiments, analyzed data, supervised the project and wrote the manuscript. E.M.T. conceived the project, secured funding, supervised the project, and wrote the manuscript. All authors approved the manuscript.

