## Supplementary Figures S1-S4 for "Reversible expression of a cartilage ‘master regulator’ during axolotl limb development and regeneration"


### Figure S1

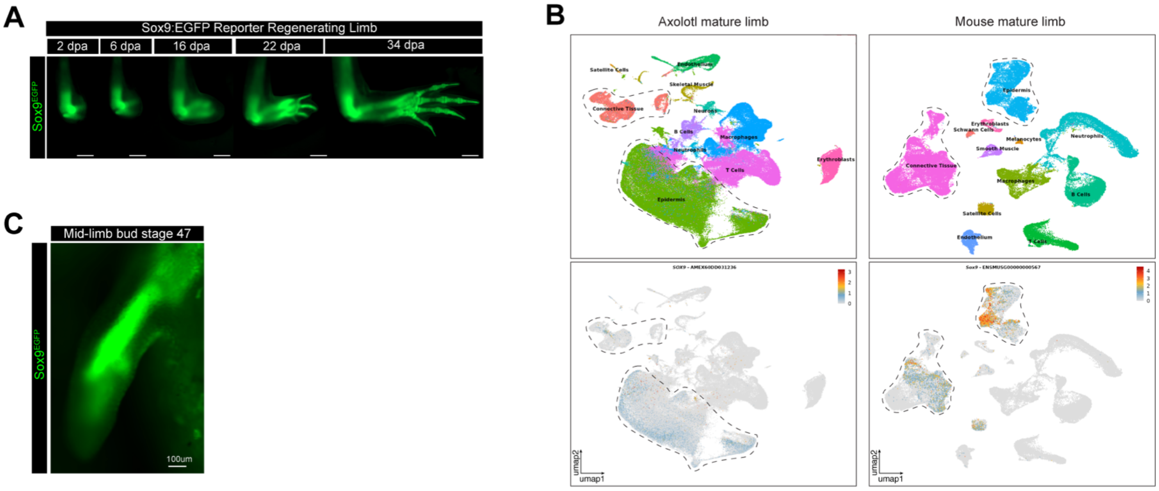


**Figure S1 Characterization of *Sox9*:EGFP-nls-T2A-ER-Cre-ER and *Sox9* expression.**

**A** Widefield images of *Sox9*:EGFP reporter during limb regeneration (*n* = 8 limbs, from 4 animals). dpa: days post-amputation.

**B** Left: UMAP representation of axolotl mature limb single cell RNA-seq data (McMann et al., 2025) showing *Sox9* expression in connective tissue (mesenchymal) and epidermis clusters. Right: UMAP representation of mouse limb single cell RNA-seq data (McMann et al., 2025) showing *Sox9* expression in connective tissue (mesenchymal) and epidermis clusters.

**C** Widefield image of *Sox9*:EGFP reporter expression at limb bud stage 47 (mid-limb bud) (Representative of *n* = 4 limb buds, from 4 animals).

### Figure S2

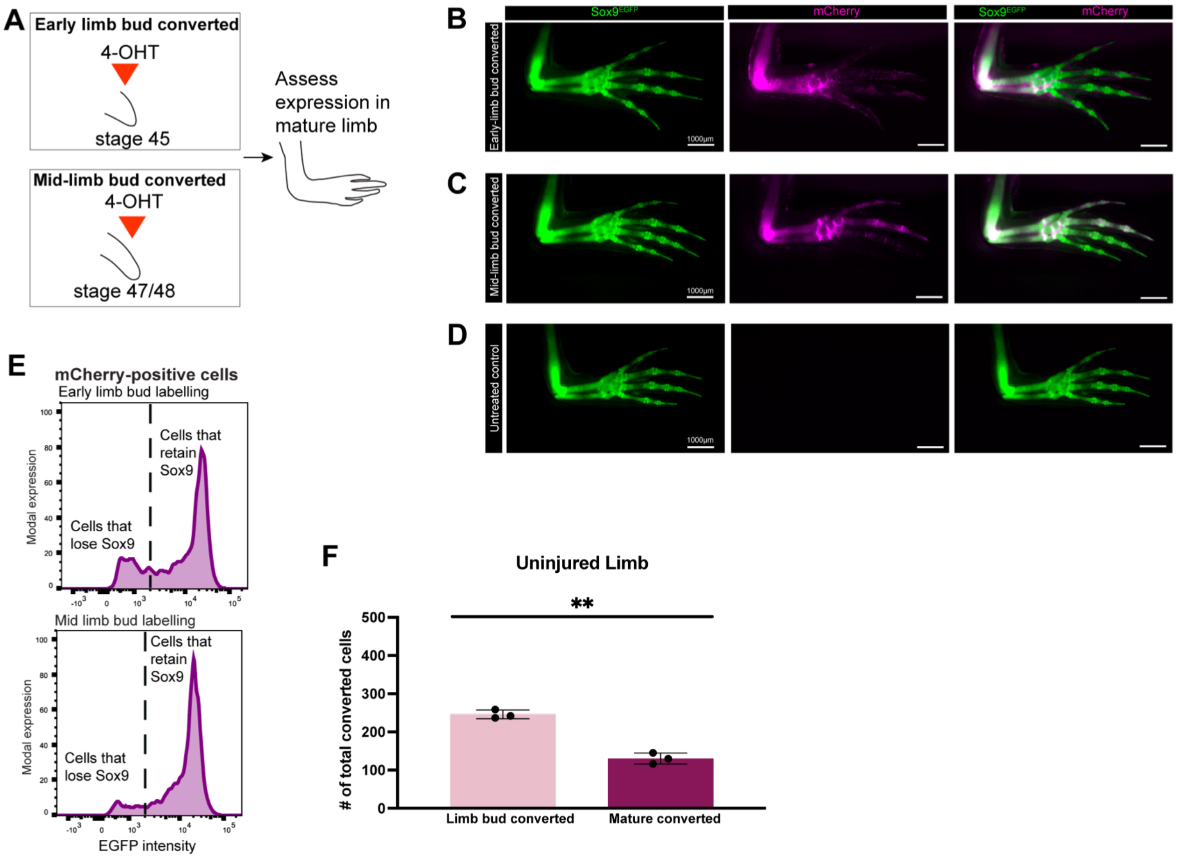


**Figure S2 Progressive restriction of *Sox9*:EGFP-nls-T2A-ER-Cre-ER expression in the developing axolotl limb**

**A** Schematics indicating the timing of conversion in genetic fate mapping experiments. After treatment with 4-OHT, animals were raised to 4 cm.

**B** Widefield images of early limb bud converted limbs (*n* = 5 animals, 2 limbs each).

**C** Widefield images of mid limb bud converted limbs (*n* = 5 animals, 2 limbs each).

**D** Widefield images of untreated control limbs (*n* = 7 animals, 2 limbs each).

**E** Flow cytometry analysis of early limb bud converted limbs (upper panel) or late limb bud converted limbs (lower panel). Panels depict EGFP fluorescence intensity in mCherry-positive cells (*n* = 4 limbs, from 2 animals per condition).

**F** Total number of converted cells per section following conversion at mature or limb bud (early and late double converted). Each dot represents one limb; bars show mean ± SD. The difference was statistically significant (two-tailed unpaired Welch's t-test on $log_{10}$-transformed values; P = 0.0043, t = 9.234, df = 2.656; *n* = 3 limbs per condition).

### Figure S3

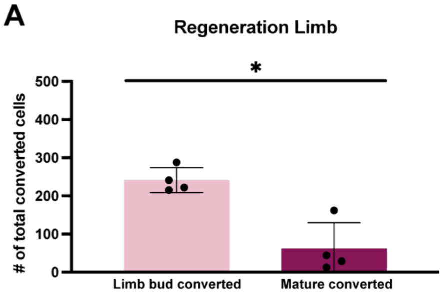


**Figure S3 Mature or limb bud converted mCherry cells**

**A** Total number of converted cells per section following conversion at mature or limb bud (early and late double converted). Each dot represents one limb; bars show mean ± SD. The difference was statistically significant (two-tailed unpaired Welch's t-test on $log_{10}$-transformed values; P = 0.0430, t = 3.325, df = 3.090; *n* = 4 limbs per condition)

### Figure S4

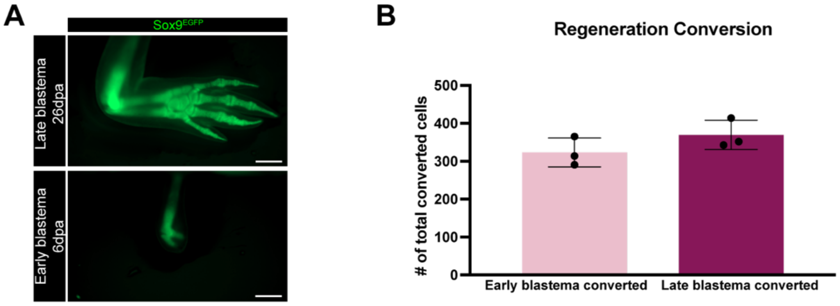


**Figure S4 Early or late blastema converted mCherry cells**

**A** Widefield image of regenerating limbs at the late (upper panel) or early (lower panel) blastema converted time points.

**B** Total number of converted cells per section following conversion at early or late blastema stages. Each dot represents one limb; bars show mean ± SD. The difference was not statistically significant (two-tailed unpaired Welch's t-test on $log_{10}$-transformed values; P = 0.2079, t = 1.505, df = 3.933; *n* = 3 limbs per condition)
